## Supplementary Figures and Tables for "Deep-learning deconvolution and segmentation of fluorescent membranes for high-precision bacterial cell-size profiling"

**Content**

- Supplementary Figures S1 to S15
- Supplementary tables S1 to S3
- Detailed descriptions of plasmid construction
- Relevant DNA sequences

SUPPLEMENTARY FIGURES

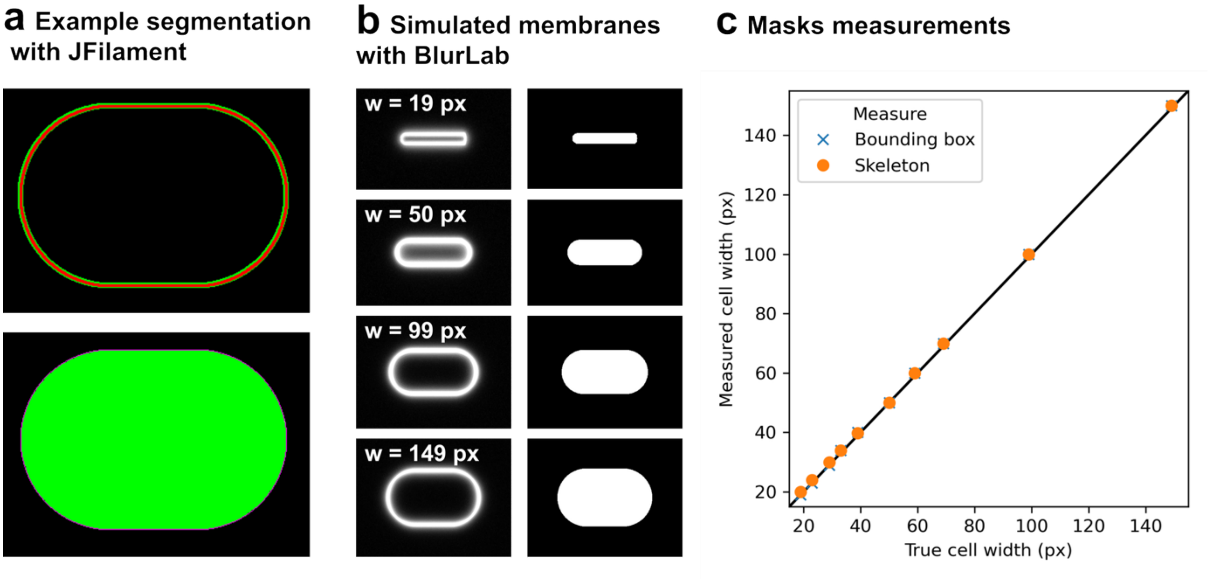


**Supplementary Figure 1. JFilament snakes accurately delineate membrane position. a**, Example of JFilament segmentation of a synthetic contour. JFilament snakes automatically adjust to and track the pixels of maximal intensity along a contour, generating a mask corresponding to the pixels enclosed by the contour. The top panel shows a JFilament snake (red) tracking a contour. The bottom panel shows the original contour (magenta) and the mask generated by JFilament (green). The mask excludes the contour pixel itself. **b**, Simulated rods of known sizes were used to generate fluorescent membrane signals with BlurLab (left) (Ursell et al., 2017), and the images were segmented with JFilament (right, JFilament masks). **c**, Rod widths measured from JFilament masks (y axis) versus true rod with (x axis). Width was estimated both by direct measurement of the mask (blue crosses; measure bounding box) and by applying the mask-based size calculation pipeline based on skeletonization described later in the manuscript (orange dots, skeleton). In both cases, the estimated widths matched the true widths closely. The diagonal black line marks a perfect correspondence between measured and true widths.

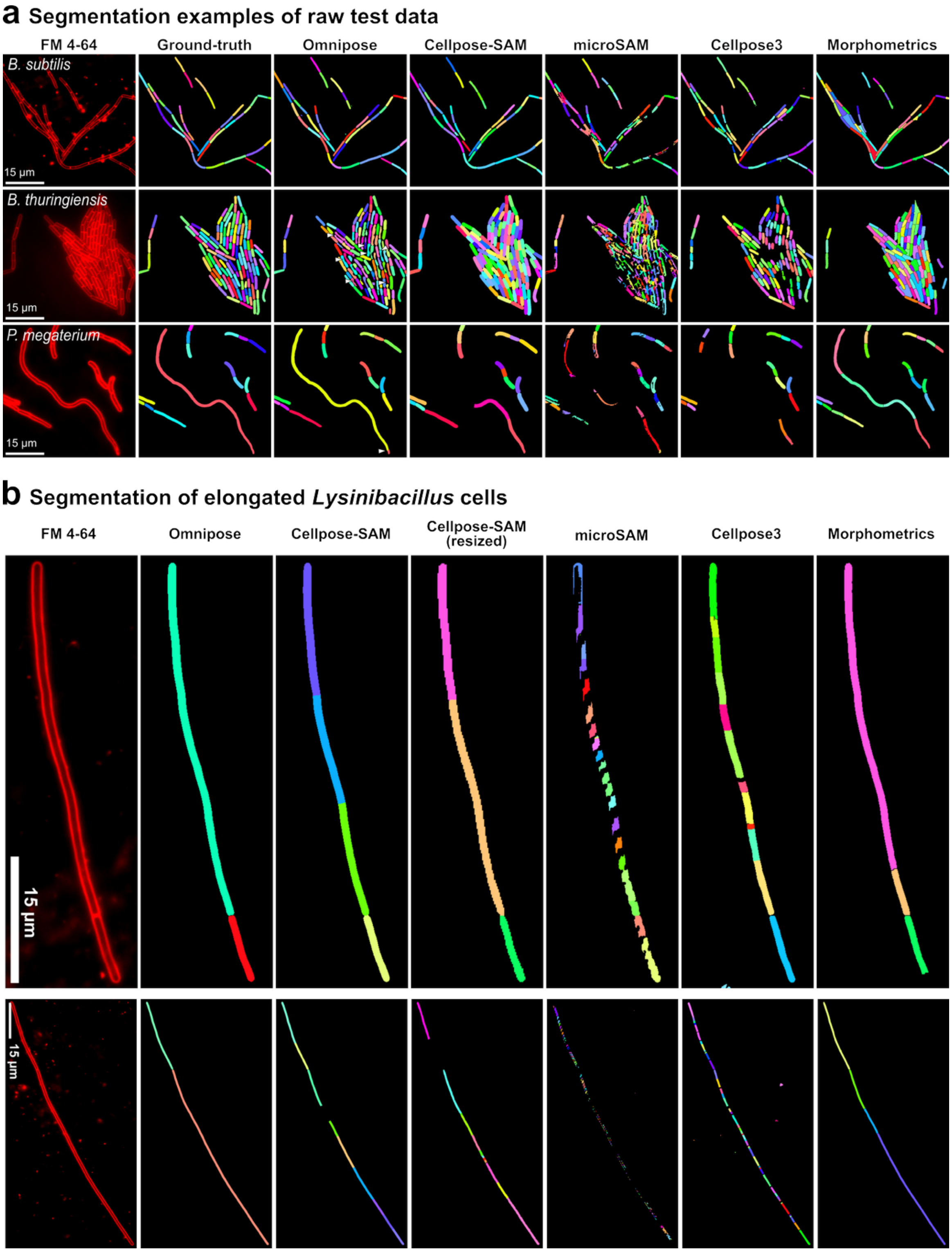


**Supplementary Figure 2. Examples of raw membrane segmentation with different algorithms.** Omnipose, Cellpose-SAM, microSAM and Cellpose3 were fine-tuned for the segmentation of non-deconvolved (raw) fluorescent membrane images. **a**, Examples of model performance on images from the benchmarking set used to calculate the F1 scores shown in Fig. 2a. First column, micrographs of *B. subtilis*, *B. thuringiensis* and *P. megaterium* cells stained with FM 4-64; second column, ground truth segmentation masks obtained with JFilament; third to seventh columns, segmentation masks obtained with the corresponding models. Masks of individual cells are in different colors. White arrowheads in the Omnipose segmentation point at representative aberrant masks in regions of clustered cells. **b**, Examples of model performance on elongated cells from a natural *Lysinibacillus* isolate. First column, micrographs of long *Lysinibacillus* cells stained with FM 4-64; second to seventh columns, segmentation results from the different models. For Cellpose-SAM, we also tried resizing the images to half the original size [Cellpose-SAM (resized)]. Scale bars, 15 µm.

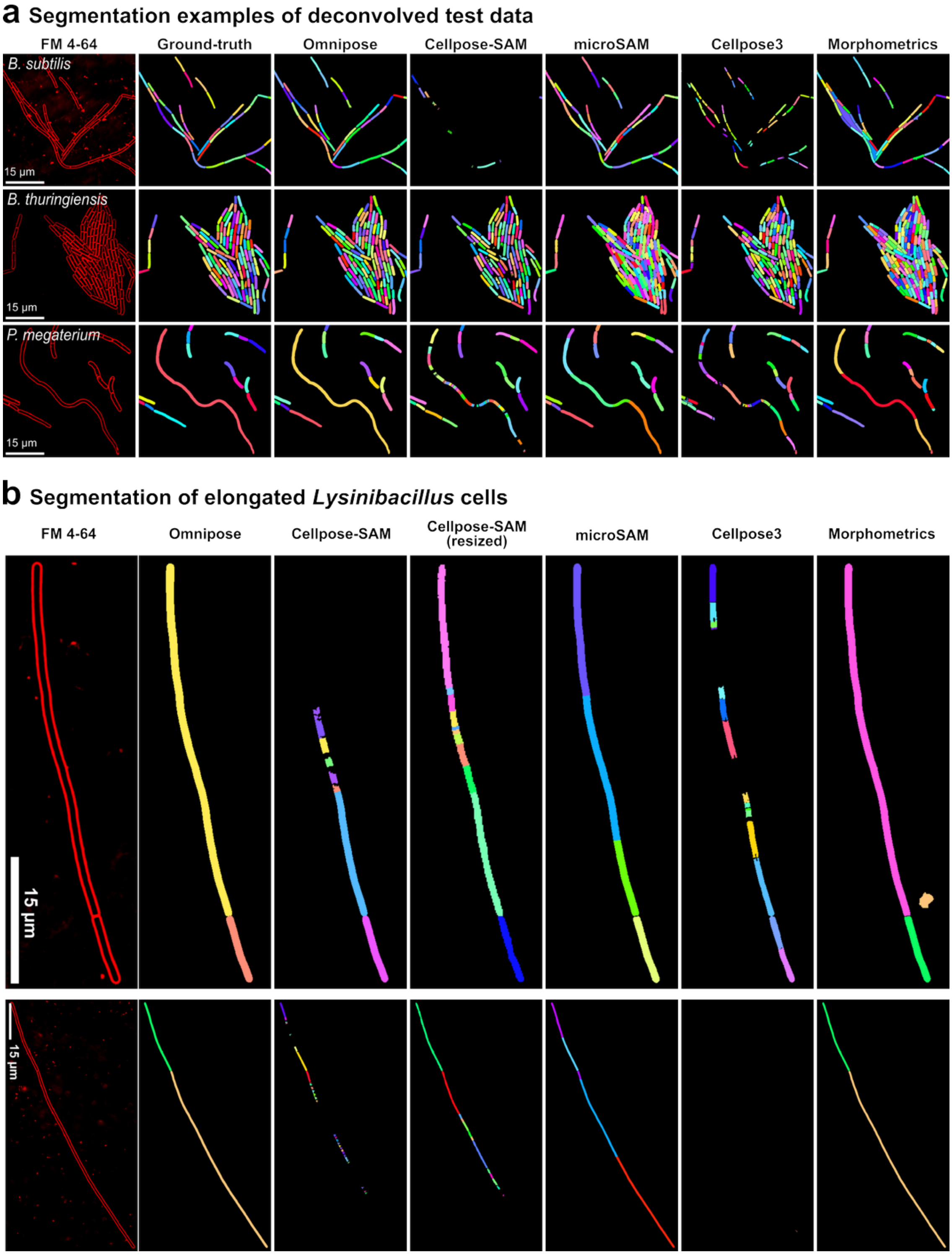


**Supplementary Figure 3. Examples of deconvolved membrane segmentation with different segmentation algorithms.** Omnipose, Cellpose-SAM, microSAM and Cellpose3 were fine-tuned for the segmentation of deconvolved fluorescent-membrane images. **a**, Examples of model performance on images from the benchmarking set used to calculate the F1 scores shown in Fig. 2a. First column, deconvolved micrographs of *B. subtilis*, *B. thuringiensis* and *P. megaterium* cells stained with FM 4-64; second column, ground truth segmentation masks obtained with JFilament; third to seventh columns, segmentation masks obtained with the corresponding models. Masks of individual cells are in different colors. **b**, Examples of model performance on elongated cells from a natural *Lysinibacillus* isolate. First column, deconvolved micrographs of long *Lysinibacillus* cells stained with FM 4-64; second to seventh columns, segmentation results from the different models. For Cellpose-SAM, we also tried resizing the images to half the original size [Cellpose-SAM (resized)]. Scale bars, 15 µm.

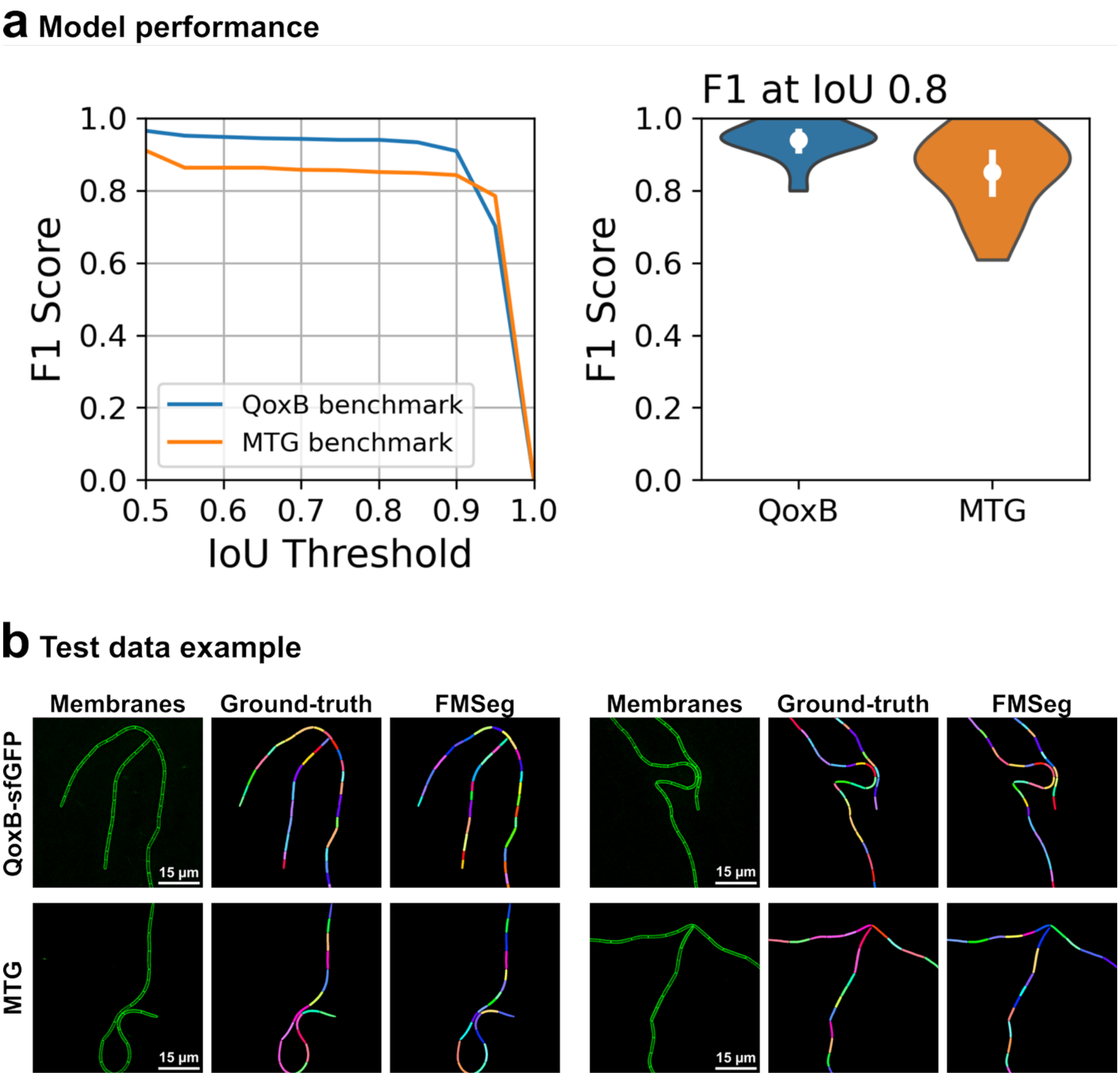


**Supplementary Fig. 4. FMSeg benchmarking on the segmentation of Mitotracker Green and QoxB-GFP fluorescence images. a**, Left, F1 score over IoU thresholds for the segmentation of membranes stained with Mitotracker green (MTG, orange) or labeled with a QoxB-GFP fusion protein (blue). The lines represent the mean F1 scored of fifteen images. Right, F1 at IoU of 0.8. The white dots and vertical lines within the violin plots represent the means and standard deviations. **B**, Examples of segmentations of QoxB-GFP and MTG images from benchmarking data. Deconvolved micrographs (membranes), ground truth masks generated with JFilament (ground truth), and FMSeg segmentation results (FMSeg) are shown. Scale bars, 15 μm.

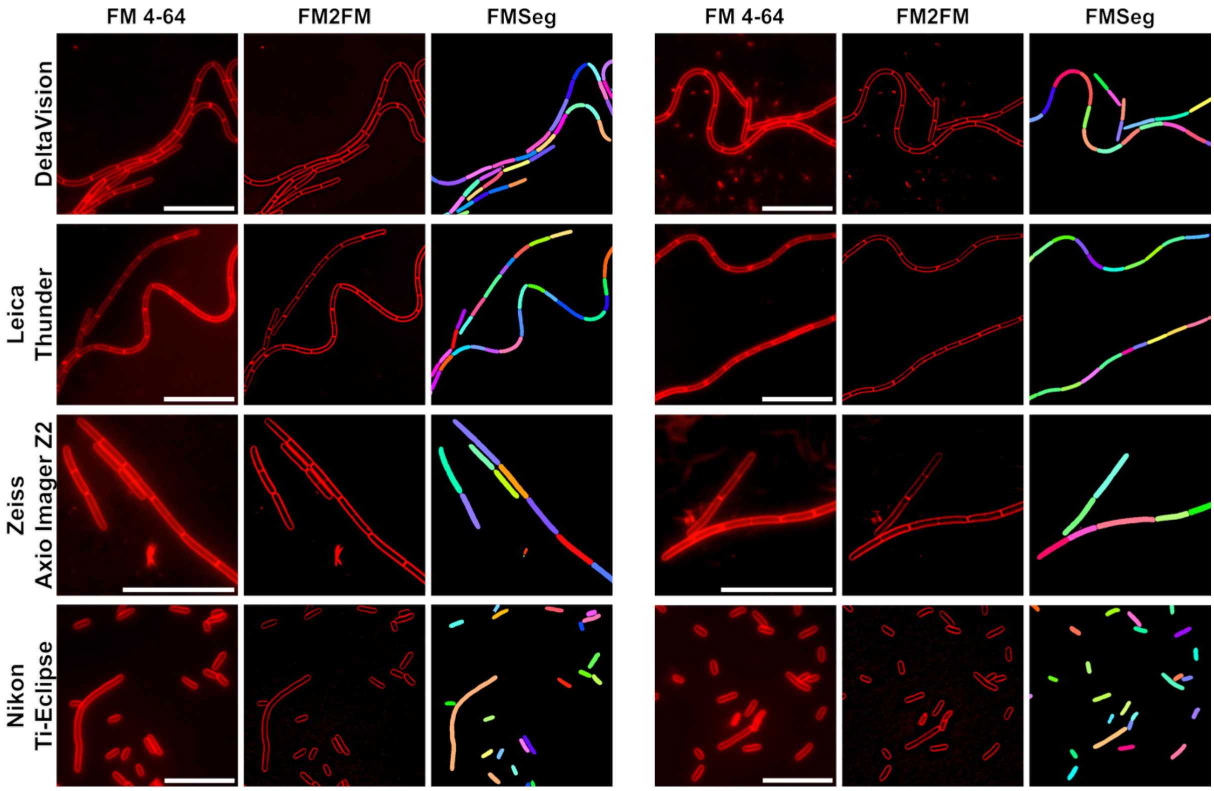


**Supplementary Fig. 5. Restoration and segmentation of images acquired with different microscopes.** Representative images of bacterial cells stained with FM 4-64, acquired using a DeltaVision, Leica Thunder, Zeiss Axio Imager Z2, or Nikon Ti-Eclipse microscope, restored with FM2FM, and segmented with FMSeg. The DeltaVision, Leica Thunder, and Zeiss Axio Imager images are from *B. subtilis* and were acquired in this study. The Nikon Ti-Eclipse images are from *E. coli* and were acquired by McKenzie and colleagues (McKenzie et al., 2022). Scale bars, 15 µm.

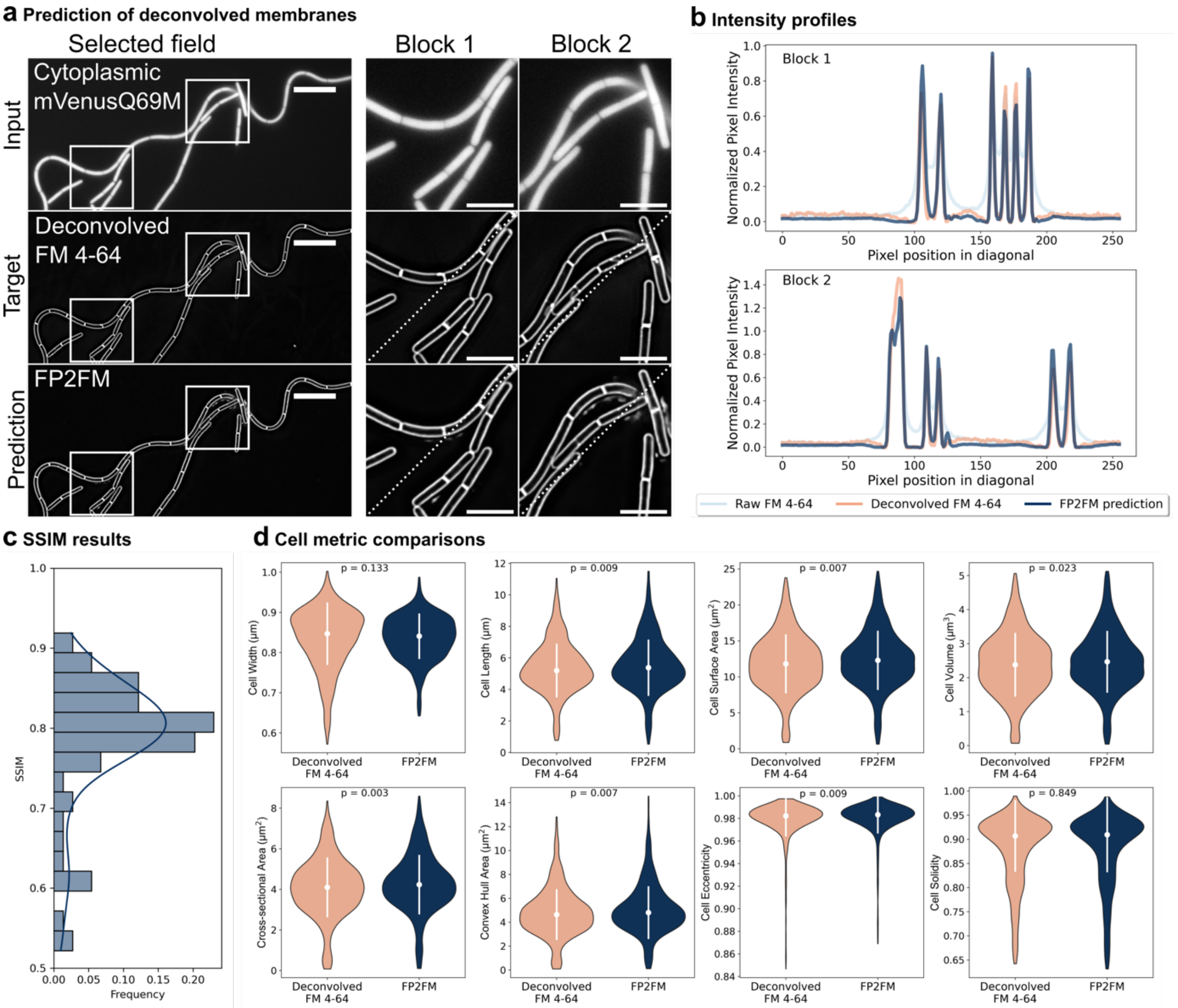


**Supplementary Fig. 6. Prediction of deconvolved membranes from cytoplasmic fluorescence.** We trained a CARE model (FP2FM) to predict deconvolved membranes from cytoplasmic fluorescence. **a**, Example of deconvolution prediction with FP2FM. Top, input raw cytoplasmic fluorescence images (Cytoplasmic mVenusQ69M); middle row, true deconvolved fluorescent membrane images (Deconvolved FM 4-64); bottom, FP2FM-predicted image. White squares mark regions zoomed in at right (Block 1 and Block 2). Diagonal dotted lines indicate profile traces in **b**. Scale bars: full images, 10 μm; zoomed-in blocks, 5 μm. **b**, Fluorescence profiles across the dotted lines in Block 1 (top) and Block 2 (bottom). Light blue, raw FM 4-64; orange, deconvolved FM 4-64; dark blue, FP2FM prediction. **c**, Structural similarity index measure (SSIM) between FP2FM predicted images and deconvolved images. The distribution of SSIM values across 74 image crops is shown. **d**, Violin plots of cell width, length, surface area, volume, cross-sectional area, convex hull area, eccentricity and solidity calculated from deconvolved FM 4-64 images (orange) and FP2FM-predicted images (dark blue). The white dots and vertical lines within the violin plots represent the medians and standard deviations. P values from statistical comparisons between distributions (ANOVA or Kruskal test; see methods for details) are indicated in each panel. Although the distributions are visually similar, statistically significant differences ($p<0.05$) were found for cell length, surface area, volume, cross-sectional area, convex hull area, and eccentricity, suggesting that deconvolved membrane prediction from cytoplasmic fluorescence could bias cell size estimations. Twenty images (over 1000 cells) were processed and segmented with FMSeg. See the Methods for size calculation details.

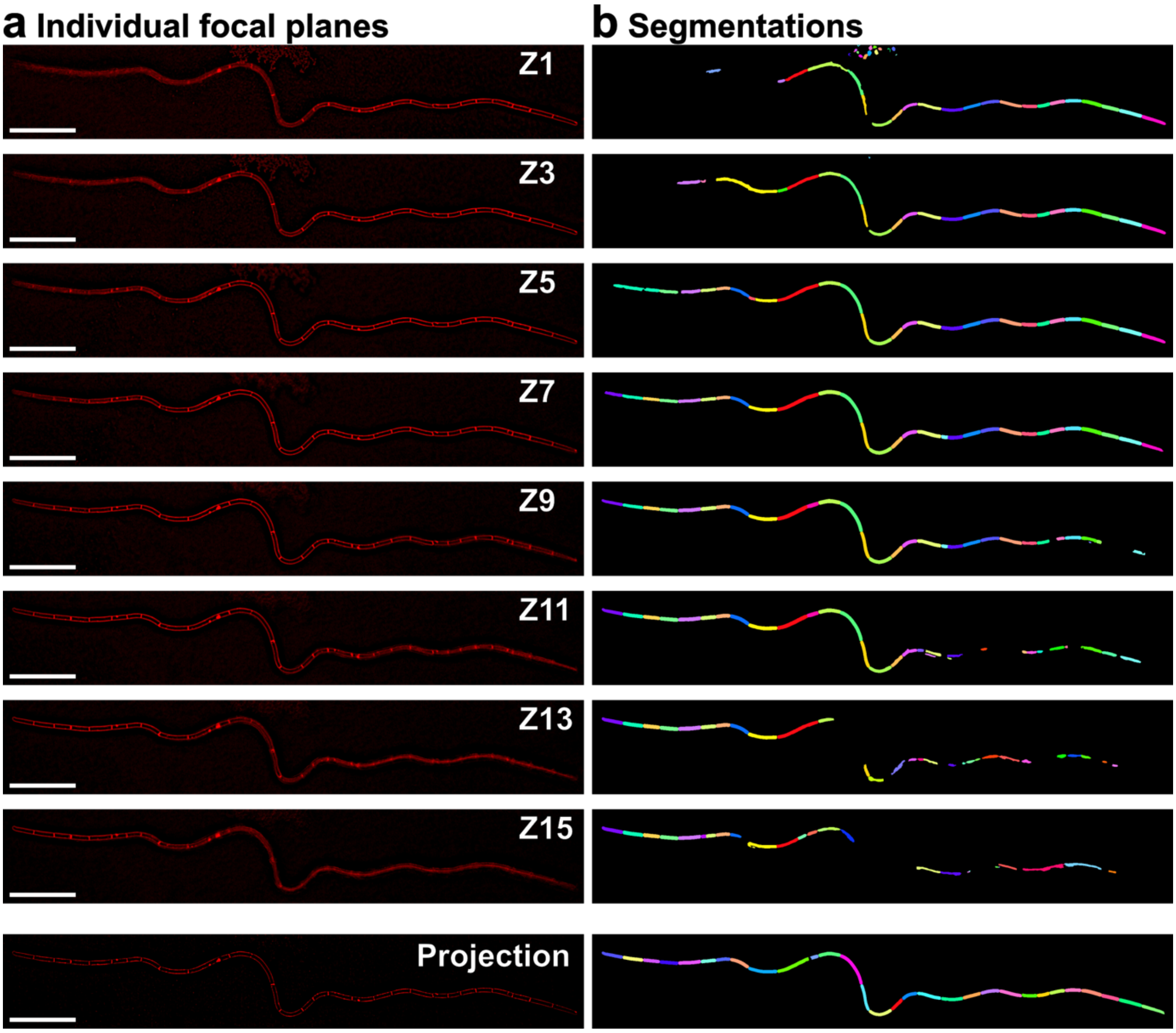


**Supplementary Figure 7. Projection and deconvolution prediction overcomes focal plane differences of individual cells in microscopy images. a**, Z-stack of chain of *B. subtilis* cells stained with FM 4-64. Individual slices are spaced 0.1 µm apart, from Z1 (top) to Z15 (bottom). The last image is a maximum intensity projection of the stack, processed using FM2FM (Projection) Scale bar = 15µm. **b**, FMSeg segmentation results of individual slices and the projected image. The projection coupled with deconvolution prediction enables segmentation of all the cells in the chain as if they were in the same focal plane.

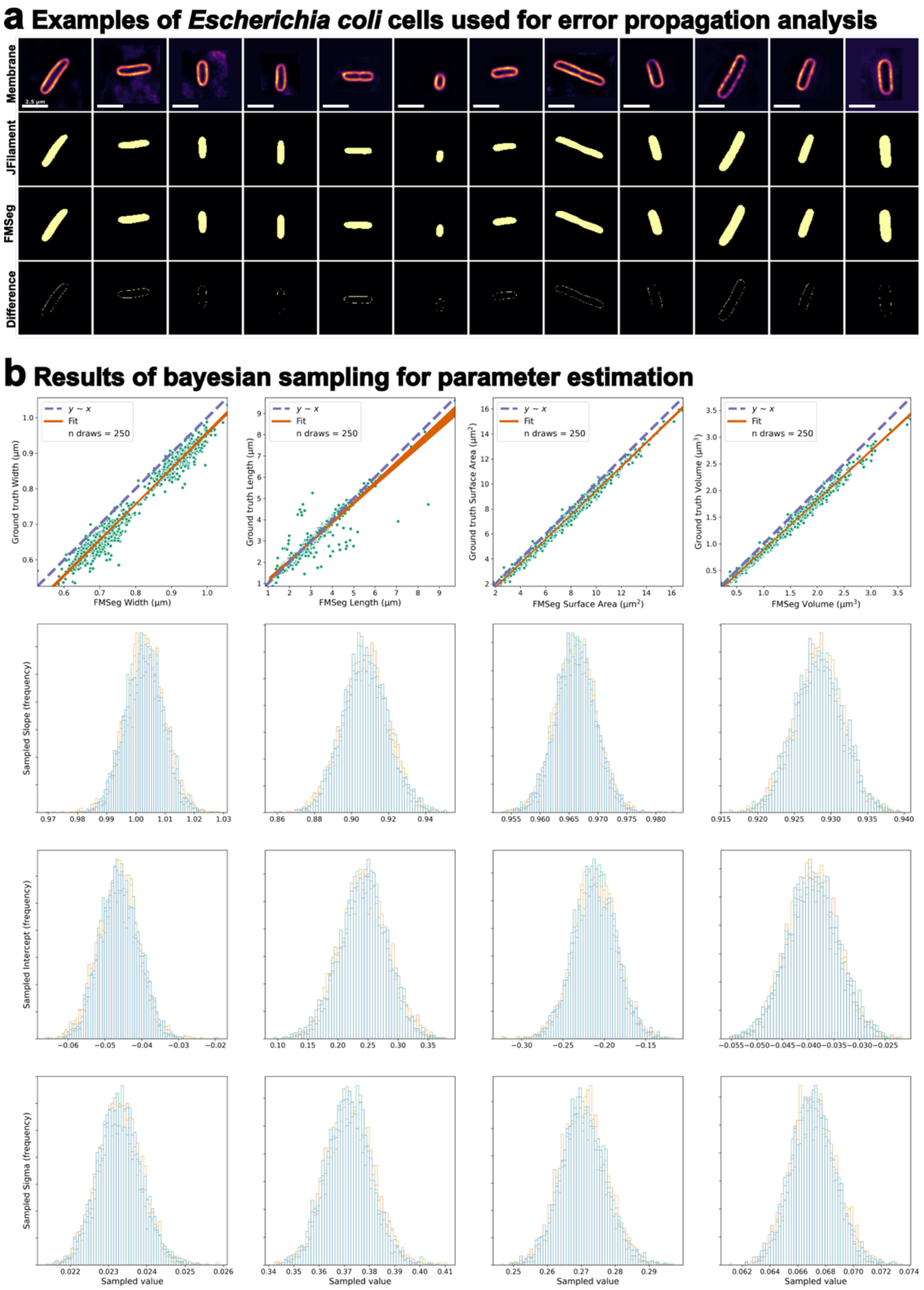


**Supplementary Figure 8. Bayesian sampling for parameter transformation.** **a**, Membrane images of *Escherichia coli* single cells (top row) were segmented both manually using JFilament (second row) and automatically using FMSeg (third row). Both segmentation approaches showed small differences in the resulting masks (bottom row). Scale bar in top row, 2.5 µm. A total of 864 cells were segmented with both approaches to quantify size differences. **b**, FMSeg produces masks slightly larger than JFilament. The scatter plots show the width, length, surface area and volume of the 864 *E. coli* cells calculated from masks generated using JFilament (ground truth value, y-axis) or automatically-generated FMSeg masks (FMSeg value, x-axis). Each green dot represents a single cell. The purple dashed line indicates a one-to-one relationship. The orange lines show 250 randomly-sampled parameter pairs used to correct size differences between FMSeg and Ground truth values. The histograms represent the drawn values for each parameter (from the second to last row: slope, intercept, and standard deviation of a re-calculated population mean). Three thousand values were sampled in four separate chains (represented by different colors in the histograms).

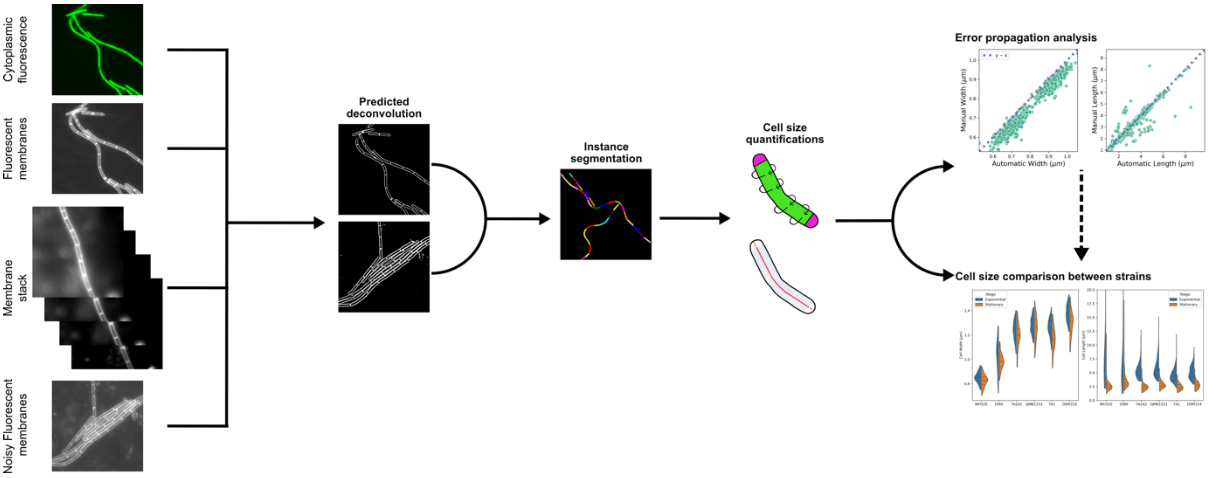


**Supplementary Fig. 9. Overview of the MEDUSSA pipeline**. MEDUSSA predicts deconvolved fluorescent membrane images that are amenable to high-quality segmentation using custom CARE models. Starting from raw fluorescent membrane images (FM2FM), cytoplasmic fluorescence (FP2FM), or low signal-to-noise raw membrane images (FM2FM-HiSNR), the pipeline restores membrane signals to a common deconvolution target. MEDUSSA also corrects focal-plane differences between cells within the same field of view by projecting z-stacks and processing the projection with FM2FM. The restored membrane images are then segmented with the fine-tuned FMSeg_Omni model to obtain instance segmentations of individual cells. Finally, MEDUSSA quantifies cell-size features and supports direct morphological comparisons across strains, including error-propagation analysis that can be used to transform the resulting measurements.

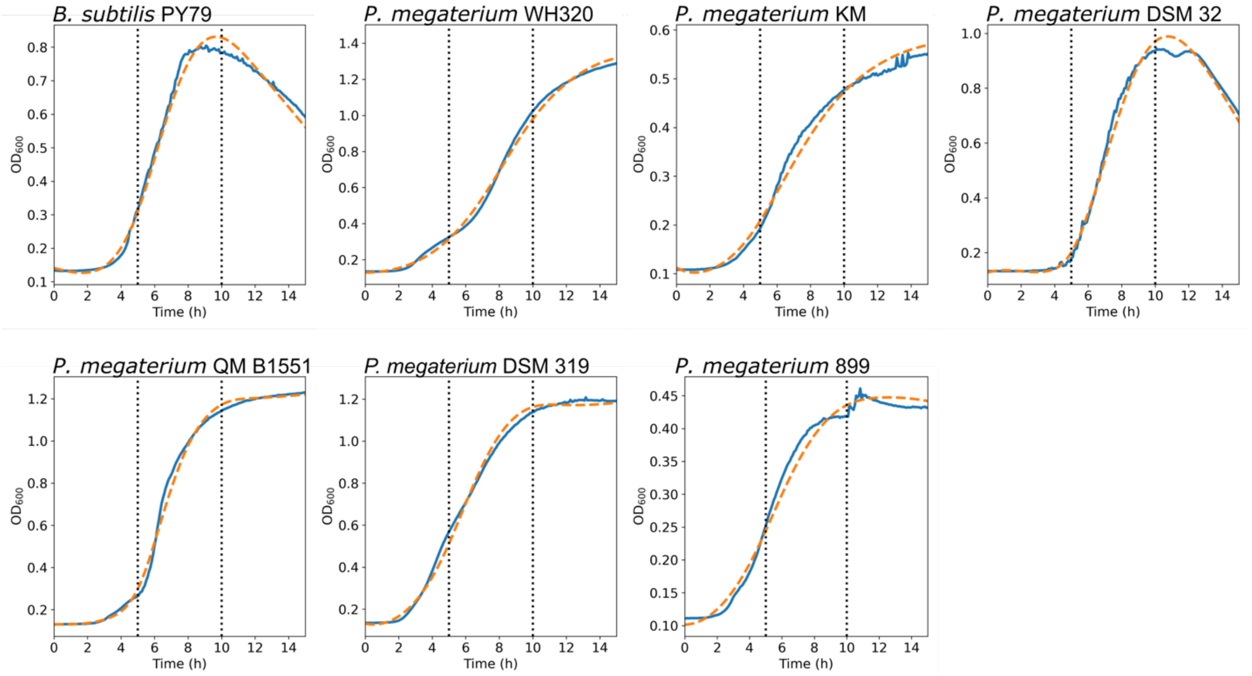


**Supplementary Figure 10. Growth curves of *P. megaterium* and *B. subtilis* strains**. Line plots show time (x-axis) versus $OD_{600}$ (y-axis) for populations of six *P. megaterium* strains (WH320, DSM 32, KM, QM B1551, DSM 319, and 899) a *B. subtilis* strain (PY79). Blue continuous line is the mean $OD_{600}$ measured across three replicates, and dashed orange line is a fitted spline curve to the mean growth data. Vertical dotted lines mark five and ten hours of growth.

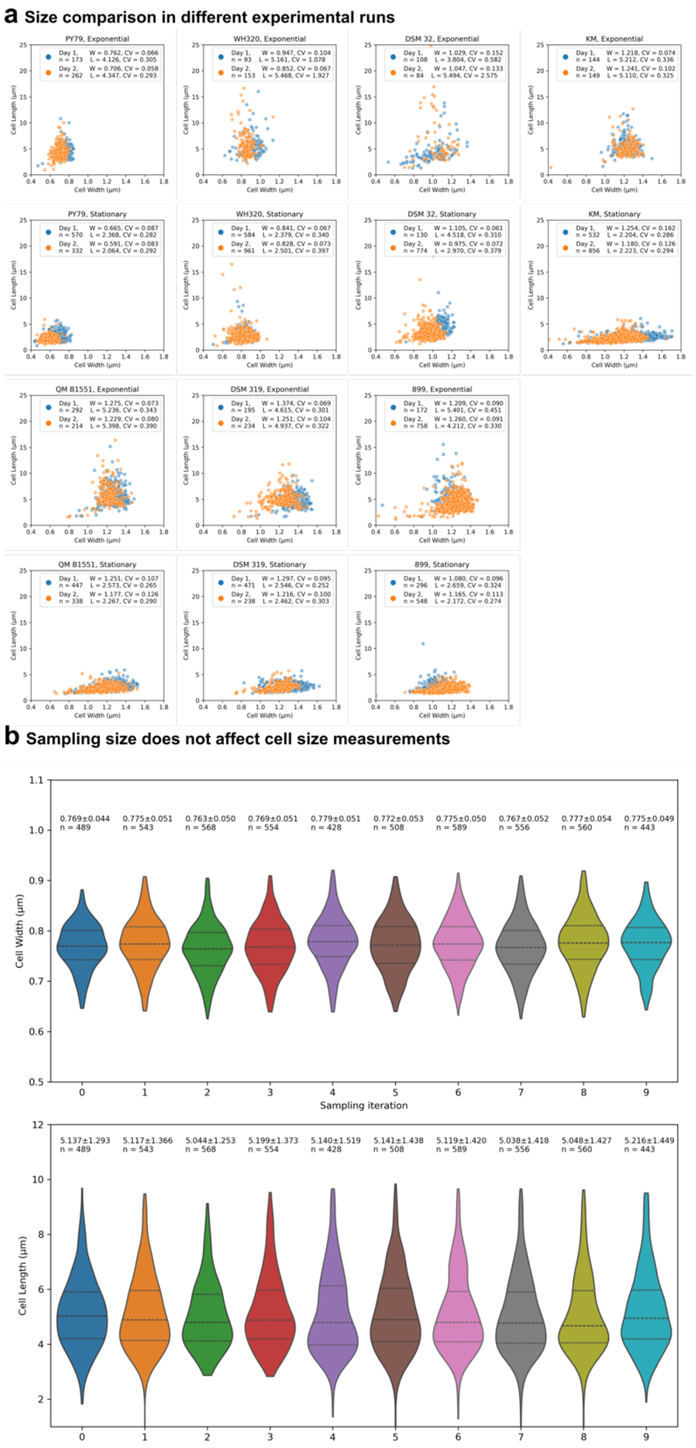


**Supplementary Figure 11. Length-width comparisons across experimental runs**. **a**, Scatter plots show cell width (x-axis) versus length (y-axis) for individual cells from the six *P. megaterium* strains in Figure 5 (WH320, DSM 32, KM, QM B1551, DSM 319, and 899) and a *B. subtilis* strain (PY79). Each dot represents a cell, colored by experimental run (blue, Day 1; orange, Day 2). Exponential- and stationary-phase distributions are shown in separate panels for each strain. The inset in each panel reports the number of cells analyzed (n), the median width (W) and length (L), and their coefficients of variation (CV). Prior to measurement, segmentation output was manually curated to remove aberrant masks and truncated masks at image edges. **b**, Width (upper graph) and length (lower graph) distributions of *B. subtilis* cells from technical replicates from the same culture. The dashed line in the violin plots indicate the median, and the dotted lines the first and third quartiles. The number of cells analyzed and the mean width or length ± standard deviation (in μm), are indicated above each violin.

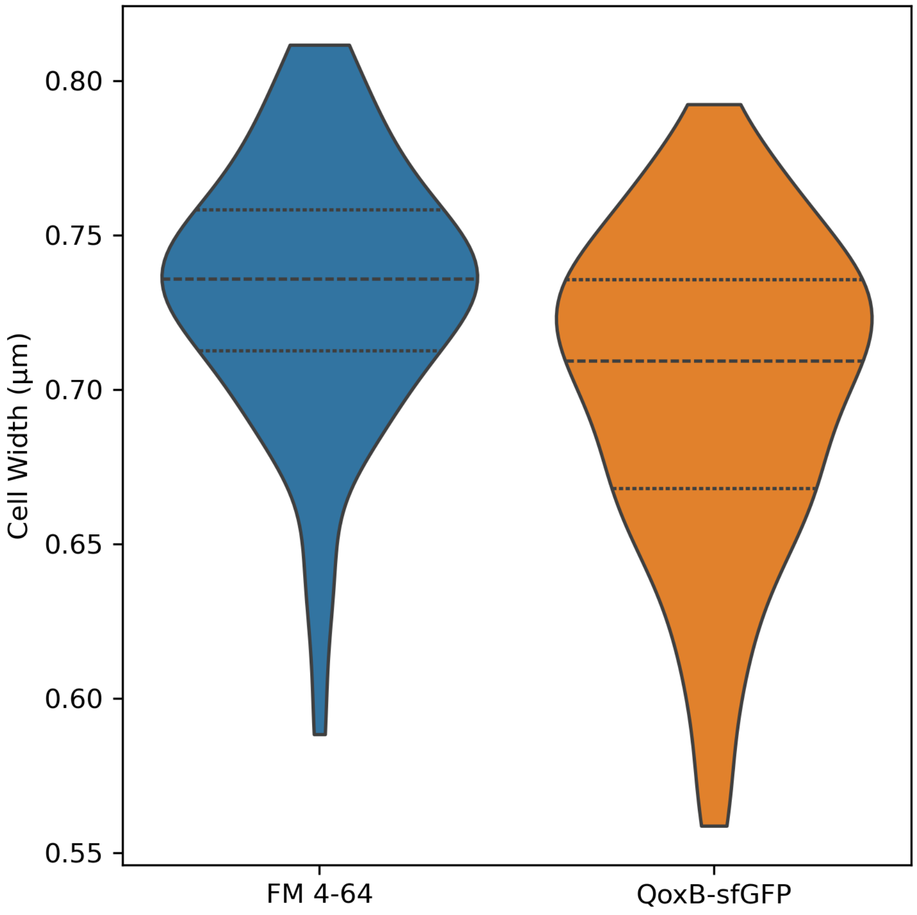


**Supplementary Fig. 12. Cell width estimation from FM 4-64 and QoxB-GFP images.** Estimates were obtained from the same field of views, in which cells carrying QoxB-GFP were stained with FM 4-64. Over 80 cells were analyzed. The dashed line in the violin plots indicate the median, and the dotted lines the first and third quartiles.

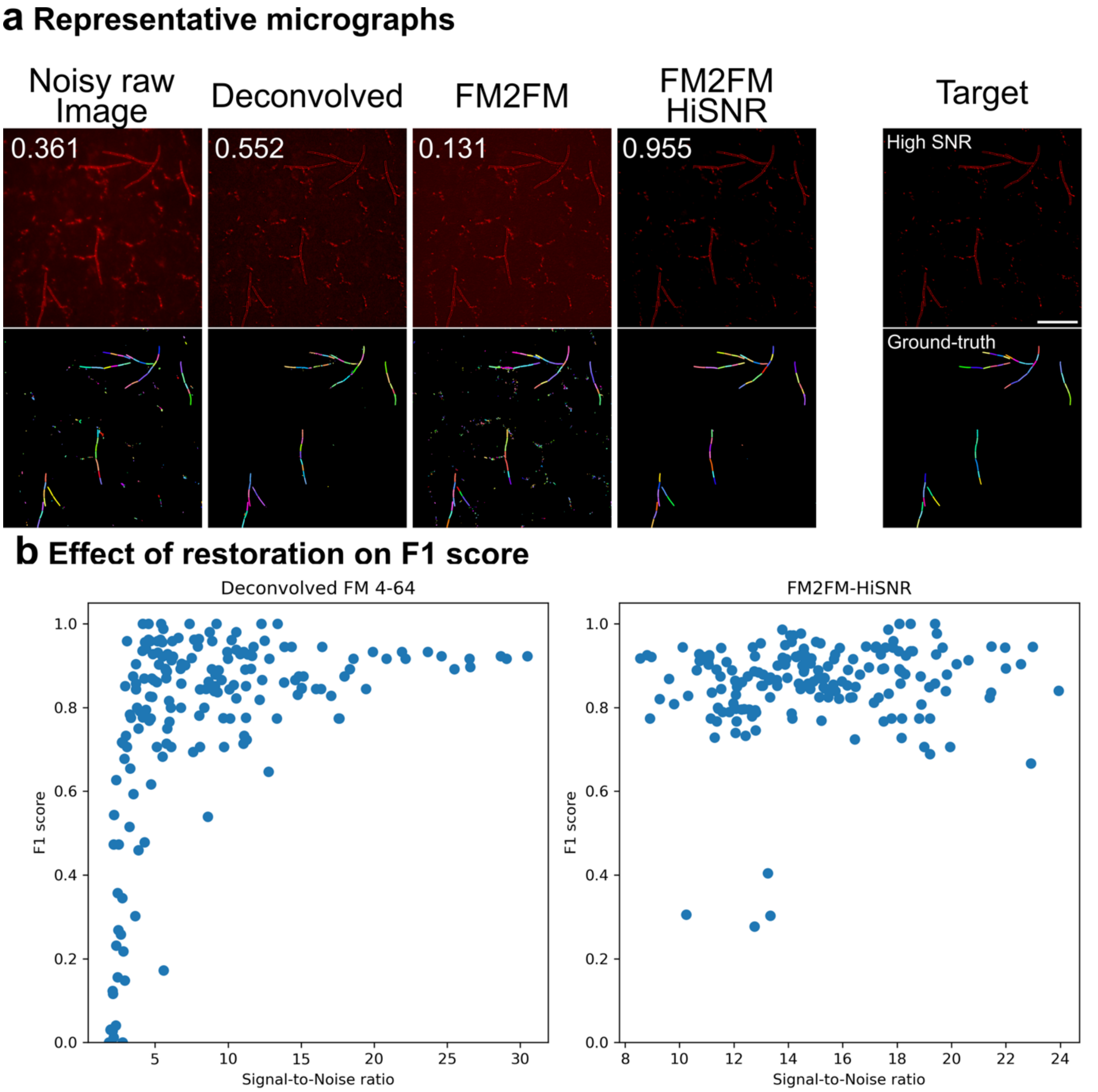


**Supplementary Fig. 13. Impact of signal-to-noise ratio on segmentation accuracy. a**, Example segmentation of a low-signal-to-noise image. Micrographs are shown at the top and the corresponding segmentation results at the bottom. First column, raw (non-deconvolved) low-signal-to-noise image (Noisy raw image) segmented with Omnipose fine-tuned for raw membrane segmentation; second column, deconvolved image segmented with Omnipose fine-tuned for deconvolved membrane segmentation (FMSeg); third column, FM2FM-predicted image segmented with FMSeg; fourth column, image predicted with the CARE model FM2FM-HiSNR, trained to predict high-signal-to-noise deconvolved images from input images spanning a range of signal-to-noise ratios, segmented with FMSeg. F1 scores at IoU 0.8 are indicated at the top-right corner of each image. The last column shows a high-signal-to-noise version of the same image, acquired at higher light exposure, which served as the target for FM2FM-HiSNR restoration and was segmented with JFilament (ground truth). **b**, F1 scores at IoU 0.8 as a function of image signal-to-noise ratio for deconvolved fluorescent membrane images (deconvolved FM4-64) and for images restored with FM2FM-HiSNR. Each blue dot represents one image (fourty images in total). FM2FM-HiSNR restoration improves segmentation quality of low signal-to-noise images. The same set of raw images was used to generate both plots.

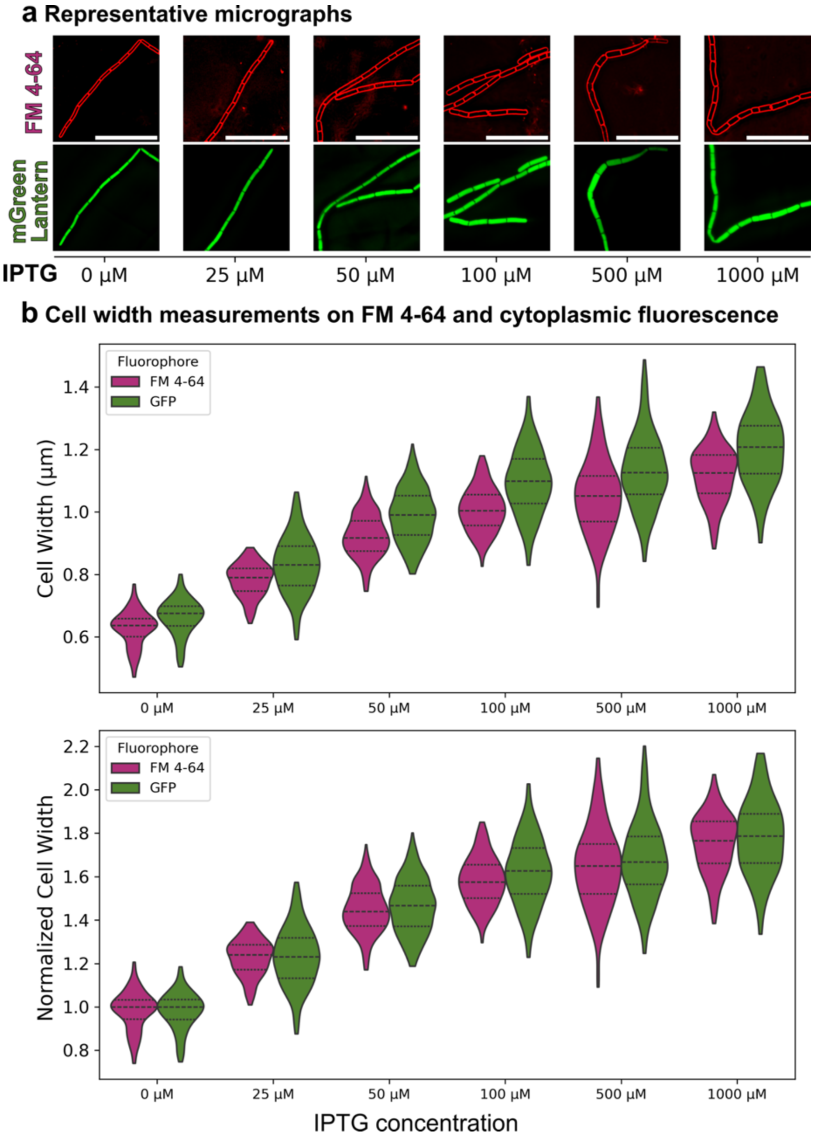


**Supplementary Fig. 14. Projection artifacts do not bias width estimates across a range of cell widths. a,** Representative micrographs of *B. subtilis* cells constitutively expressing cytoplasmic mNeonGreen and carrying the sole *ponA* copy under the IPTG-inducible Pspank promoter, grown at different IPTG concentrations. In the absence of IPTG, cells are thin, similar to a *ponA* mutant, whereas increasing IPTG progressively increases average cell width. Deconvolved FM 4-64 membrane fluorescence and cytoplasmic mNeonGreen fluorescence are shown. Scale bars, 10 µm. **b**, Cell width distributions estimated independently from membrane fluorescence (magenta; segmented with FMSeg) and cytoplasmic fluorescence (green; segmented with Cellpose-SAM) across cultures grown at different IPTG concentrations. This experiment was designed to test whether projection artifacts were affecting our measurements. Hardo et al. (Hardo et al., 2024) described projection artifacts that have opposite effects depending on the fluorescent marker: membrane fluorescence tends to underestimate true width, whereas cytoplasmic fluorescence tends to overestimate it. The magnitude of both biases was reported to be width-dependent, with proportionally larger effects in thinner cells (becoming very apparent in cells below 1 µm) than in thicker cells. If such effects were prominent under our imaging and analysis conditions, width estimates from cytoplasmic fluorescence would be expected to exceed membrane-based estimates more strongly in thin cells than in thick cells. Instead, the two measurements scaled similarly across the full width range. The upper graph shows absolute width values, and the lower graph shows normalized values, with mean widths for each segmentation mode set to 1 in the absence of IPTG. Absolute median widths were ~5–10% larger for mNeonGreen-based segmentations at all IPTG concentrations. Because masks were generated using different models for the two fluorescence signals, absolute widths are not expected to coincide exactly. However, the relative offset between the two measurements remained approximately constant across the tested width range, indicating that projection artifacts do not bias our width estimates, at least within the tested range of cell widths. At least 130 cells were measured per condition.

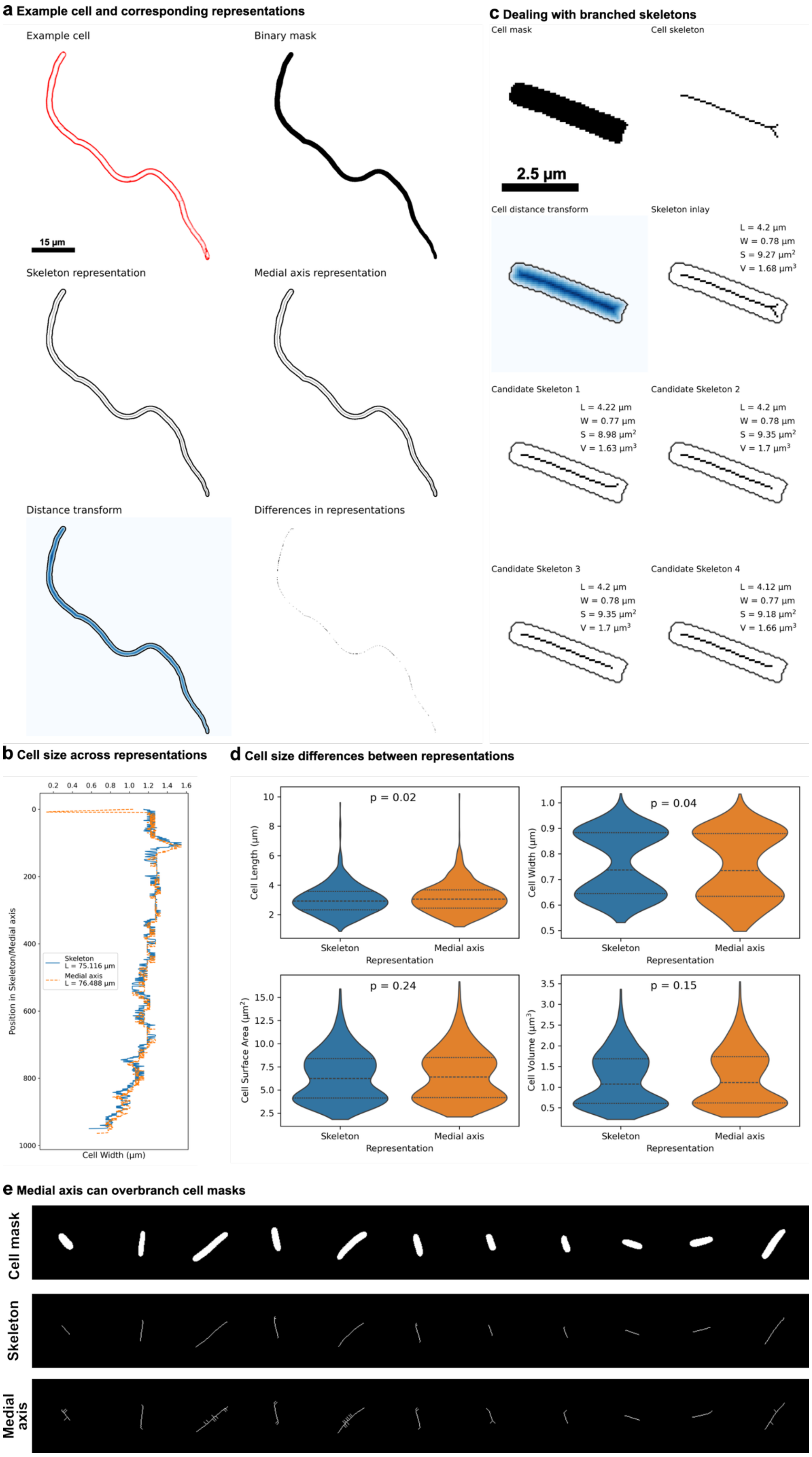
a Example cell and corresponding representations
Example cell
Binary mask
15 µm
Skeleton representation
Medial axis representation
Distance transform
Differences in representations
c Dealing with branched skeletons
Cell mask
Cell skeleton
2.5 µm
Cell distance transform
Skeleton inlay
L = 4.2 µm
W = 0.78 µm
S = 9.27 µm²
V = 1.68 µm³
Candidate Skeleton 1
L = 4.22 µm
W = 0.77 µm
S = 8.98 µm²
V = 1.63 µm³
Candidate Skeleton 2
L = 4.2 µm
W = 0.78 µm
S = 9.35 µm²
V = 1.7 µm³
Candidate Skeleton 3
L = 4.2 µm
W = 0.78 µm
S = 9.35 µm²
V = 1.7 µm³
Candidate Skeleton 4
L = 4.12 µm
W = 0.77 µm
S = 9.18 µm²
V = 1.66 µm³
b Cell size across representations
Skeleton
L = 75.116 µm
Medial axis
L = 76.488 µm
Position in Skeleton/Medial axis
Cell Width (µm)
d Cell size differences between representations
p = 0.02
p = 0.04
p = 0.24
p = 0.15
Cell Length (µm)
Cell Width (µm)
Cell Surface Area (µm²)
Cell Volume (µm³)
Skeleton
Medial axis
Representation
e Medial axis can overbranch cell masks
Cell mask
Skeleton
Medial axis

**Supplementary Figure 15. Details of the imaging pipeline.** **a**, Comparison of skeleton and medial-axis representations. Both were computed from a binary mask of an example FM 4-64-stained cell; pixel-wise differences between the skeleton and medial axis are shown in the bottom right panel. Scale bar, 15 µm. **b**, Comparison of the local widths and total cell length calculated using the skeleton (blue) or the medial axis (orange). Although similar, the two representations show slight differences in their local values. **c**, Graph strategy for branching skeletons. Four candidate skeletons were generated and size metrics computed for each; a single value per metric was then taken as the median across candidates (values shown inset in the "Skeleton inlay" panel). Scale bar, 2.5 µm. **d**, Cell size quantification of *E. coli* cells (ground truth masks used in Supplementary Fig. 8). Blue violins are the values obtained with the skeleton and orange violins are the values obtained with the medial axis. P values are indicated between violins (obtained from either an ANOVA or Kruskal test, see methods for details). **e**, Skeleton and medial axis representations of representative cells. Medial axis representation tended to over-branch more than the skeleton representation, which might explain the differences in some of the estimated cell size parameters. We proceeded with the skeleton representation.

**Supplementary Table 1.** Strains used in this study.

| **Strain** | **Genotype** | **Source/Construction[a]** |
|---|---|---|
| PY79 | Wild-type | (Youngman et al., 1984) |
| JLG256 | Wild-type *B. subtilis* 168 | |
| JLG1561 | Δ*ponA*::Km | (Tocheva et al., 2013) |
| JLG2663 | *rpsB-sfGFP*ΩEm | pJLG396 → PY79 |
| JLG3576 | *qoxB-sfGFP-loxP*-Em-*loxP* | pJLG650 → PY79 (Erm) |
| JLG3974 | *yxiA*::$P_c$-*mNeonGreen*ΩKm | (Kasu et al., 2024) |
| JLG3976 | *yxiA*::$P_c$-*mVenusQ69M*ΩKm | pJLG912 → PY79 (Km) |
| JLG6822 | Δ*ponA*::Km *amyE*::$P_{spank}$-*ponA*$_{WH320}$ΩSp | pJLG1428 → JLG1561 (Sp) |
| JLG6827 | Δ*ponA*::Km *amyE*::$P_{spank}$-*ponA*$_{PY79}$ΩSp | pJLG929 → JLG1561 (Sp) |
| JLG6872 | Δ*ponA*::Km *amyE*::$P_{spank}$-*ponA*$_{DSM319}$ΩSp | pJLG1440 → JLG1561 (Sp) |
| JLG7125 | Δ*ponA*::Km *amyE*::$P_{spank}$-*ponA*$_{PY79}$ΩSp *yxiA*::$P_c$-*mGreenLantern*ΩErm | pJLG1307 → JLG6827 (Erm) |
| DSM 32 | *P. megaterium* DSM 32 | Leibniz Institute DSMZ |
| DSM 319 | *P. megaterium* DSM 319 | Leibniz Institute DSMZ |
| JLG2644 | *P. megaterium* WH320 | MoBiTec |
| JLG2645 | *P. megaterium* QM B1551 | Dr. Peter Setlow at UConn Health |
| BGSC 7A1 | *P. megaterium* 899 | Bacillus Genetic Stock Center |
| BGSC 7A2424 | *P. megaterium* KM | Bacillus Genetic Stock Center |
| DSM 11869 | *Lactococcus lactis* SB 150 | Leibniz Institute DSMZ |
| CECT11094 | *Saccharomyces cerevisiae* S288C | Colección Española de Cultivos Tipo |
| JLG2647 | *Bacillus pumilus* BL8 | Dr. Louise Temple at James Madison University |
| CGSC8237 | *Escherichia coli* MG1655 | Coli Genetic Stock Center |
| BGSC 13A2 | *Lysinibacillus sphaericus* SS II-1 | *Bacillus* Genetic Stock Center |
| BGSC 4AF1 | *Bacillus thuringiensis serovar cameroun* T32001 | *Bacillus* Genetic Stock Center |
| BGSC 4Q3 | *Bacillus thuringiensis serovar israelensis* IPS 70 | *Bacillus* Genetic Stock Center |
| V000017 | *Micrococcus sp.* | This study[b] |

[a]Plasmids shown to the left of the arrow were used to transform the strains on the right to generate the corresponding strain. The antibiotic used for selection is given in parentheses.

[b]Isolated from the soil

**Supplementary Table 2.** Plasmids used in this study.

| **Plasmid** | **Description** | **Reference** |
|---|---|---|
| pDG1662 | *amyE*::Cm | (Guérout-Fleury et al., 1996) |
| pDG1731 | *thrC*::Sp | (Guérout-Fleury et al., 1996) |
| pDR110 | *amyE*::($P_{spank}$, *lacI*)ΩSp | Gift from David Rudner |
| pER226 | *rpsB-GFPΩkan* | (Lopez-Garrido et al., 2018) |
| pJLG396 | *rpsB-sfGFP*ΩEm | This study |
| pJLG650 | *qoxB-sfGFP*-loxP-Erm-loxP | This study |
| pJLG884 | *yxiA*::$P_c$-*tomato*ΩKm | (Kasu et al., 2024) |
| pJLG912 | *yxiA*::$P_c$-*mVenus*$_{Q69M}$ΩKm | This study |
| pJLG929 | *amyE*::($P_{spank}$-*ponA*$_{PY79}$, *lacI*)ΩSp | This study |
| pJLG946 | *yxiA*::$P_c$-*mGreenLantern*ΩKm | This study |

| | | |
|---|---|---|
| pJLG1307 | *yxiA*::$P_c$-*mGreenLantern*ΩErm | This study |
| pJLG1428 | *amyE*::($P_{spank}$-*ponA*$_{WH320}$, *lacI*)ΩSp | This study |
| pJLG1440 | *amyE*:: ($P_{spank}$-*ponA*$_{DSM319}$, *lacI*)ΩSp | This study |

**Supplementary Table 3.** Oligonucleotides used in this study.

| **Oligo** | **Sequence**[a] |
|---|---|
| oER419 | ttctgctccctcgctcaggcggccgcTCCTTTAACTCTGGCAACCC |
| oER420 | cagggagcactggtcaacgctagcATGTGATAACTCGGCGTATG |
| oJLG7 | AATTGGGACAACTCCAGTG |
| oJLG77 | GCTAGCAGCGCAAGCGC |
| oJLG96 | GCACTTTTCGGGGAAATGTG |
| oJLG1696 | catggattacgcgttaacccAGGTCTAGAGGATCGATCTG |
| 38160_Front_F | gggttaacgcgtaatccatgCGTGAAATCAGCGGAGATTC |
| 38160_Front_R | tgcgcttgcgctgctagcTTCGGAAATCTTTCTTTCCGTC |
| 38160_Back_F | cactggagttgtcccaattTAAGGAGGCGTGAGTTATGG |
| 38160_Back_R | cacatttccccgaaaagtgcCGATTGTTACGTGAGTTCCG |
| oJLG1406 | TACCTAGATTTAGATGTCTAAAAAGC |
| oJLG1407 | TTATTATTTCCTTCCTCTTTTCTAC |
| oJLG1408 | gtagaaaagaggaaggaaataataaATGAACGAGAAAAATATAAAACACAG |
| oJLG1409 | gctttttagacatctaaatctaggtaGCAGTTTATGCATCCCTTAAC |
| oJLG1827 | TGATAAATGAGAGAGGAAGAAAAC |
| oJLG1828 | CATTCTAAAATCCTCCTTACACAC |
| oJLG1833 | gtgtgTAAGGAGGATTTTAGAATG |
| oJLG1834 | gttttcttcctctcTCATTTATCA |
| oJLG1872 | AAGCTAATTCGGTGGAAACGAG |
| oJLG1876 | CGACTAAGCTTAATTGTTATCCGC |
| oJLG2071 | gcggataacaattaagcttagtcGTTAAGGAGGAACTACCATGTCAGATCAATTTAACAGCCGTG |
| oJLG2072 | ctcgtttccaccgaattagctTTTAATTTGTTTTTTCAATGGATGATGAGTT |
| oJLG2657 | GCCTGAGCGAGGGAGCAGAActtcatcctcttcgtcttgg |
| oJLG2658 | GCGTTGACCAGTGCTCCCTGAAGTACATCCGCAACTGTCC |
| oJLG3152 | gcggataacaattaagcttagtcGCGTTGAAAGGTAGGAGTTTTTATGTC |
| oJLG3153 | ctcgtttccaccgaattagctTGCTTATTCTTCACAAGAAATAAAGCC |

[a]Primer regions that align to the template are in upper case, homology regions for Gibson assembly in lower case.

**Detailed descriptions of plasmid construction**

**pJLG396.** Two fragments were joined by Gibson isothermal assembly (New England Biolabs): (i) an inverse PCR fragment from pER226 (Lopez-Garrido et al., 2018) amplified with primers oJLG1406 and oLG1407; (ii) the coding sequence of erythromycin resistance gene from pDG1731 (Guérout-Fleury et al., 1996), amplified with oJLG1408 and oJLG1409. The kanamycin resistance gene from pER226 is replaced by the erythromycin-resistance gene from pDG1731.

**pJLG650.** Four DNA fragments were joined by Gibson isothermal assembly (New England Biolabs): (i) a 384-bp fragment corresponding to the 3' end of *qoxB* coding sequence without the stop codon, amplified with oligos 38160_Front_F and 38160_Front_R; (ii) *sfGFP*-loxP-Erm-loxP fragment, amplified with oligos oJLG7 and oJLG77 from genomic DNA of strain JLG2663; (iii) a 467-bp corresponding including the stop codon of *qoxB* and the region immediately downstream of it, amplified with oligos 38160_Back_F and 38160_Back_R); (iv) a DNA fragment encompassing the spectinomycin resistance gene, the origin of replication, and the ampicillin resistance gene from pDG1662 (Guérout-Fleury et al., 1996), amplified with primers oJLG96 and oJLG1696.

**pJLG912.** Two fragments were joined by Gibson isothermal assembly (New England Biolabs): (i) a fragment of 735 bp containing the coding sequence of mVenusQ69M (Cox et al., 2010), generated by gene synthesis (Biomatik; sequence provided below below) and amplified with primers oJLG1833 and oJLG1834; (ii) an inverse PCR product of pJLG884 that leaves out the tomato coding sequence generated by amplification with primers oJLG1827 and oJLG1828. In the resulting plasmid, the tomato coding sequence of pJLG884 is replaced by mVenusQ69M.

**pJLG929.** Two fragments were joined by Gibson isothermal assembly (New England Biolabs): (i) a fragment of 2786 bp containing the RBS, coding sequence, and putatitve terminator of *B. subtilis* PY79 *ponA*, amplified from genomic DNA of *B. subtilis* PY79 with primers oJLG2071 and oJLG2072; (ii) an inverse PCR product of pDR110 to introduce the amplified sequence downstream of the Pspank promoter, amplified with primers oJLG1872 and oJLG1876.

**pJLG946.** Two fragments were joined by Gibson isothermal assembly (New England Biolabs): (i) a fragment of XX bp containing the coding sequence of mGreenLantern, generated by gene synthesis (Biomatik; sequence provided below below) and amplified with primers oJLG1833 and oJLG1834; (ii) an inverse PCR product of pJLG884 that leaves out the tomato coding sequence generated by amplification with primers oJLG1827 and oJLG1828. In the resulting plasmid, the tomato coding sequence of pJLG884 is replaced by mGreenLantern.

**pJLG1307.** Two fragments were joined by Gibson isothermal assembly (New England Biolabs): (i) a fragment of 1395 bp containing an antibiotic resistance cassette of Erythromycin amplified with primers oER419 and oER420 from pDG1731 [7]; (ii) an inverse PCR product of pJLG946 that leaves out the Kanamycin resistance cassette generated by amplification with primers oJLG2657 and oJLG2658.

**pJLG1428.** Two fragments were joined by Gibson isothermal assembly (New England Biolabs): (i) a fragment of 2927 bp containing the RBS, coding sequence, and putatitve terminator of *P. megaterium* WH320 *ponA*, amplified from genomic DNA of *P. megaterium* WH320 with primers oJLG3152 and oJLG3153; (ii) an inverse PCR product of pDR110 to introduce the amplified sequence downstream of the Pspank promoter, amplified with primers oJLG1872 and oJLG1876.

**pJLG1440.** Two fragments were joined by Gibson isothermal assembly (New England Biolabs): (i) a fragment of 2927 bp containing the RBS, coding sequence, and putatitve terminator of DSM319 *ponA*, amplified from genomic DNA of *P. megaterium* DSM 319 with primers oJLG3152 and oJLG3153; (ii) an inverse PCR product of pDR110 to introduce the amplified sequence downstream of the Pspank promoter, amplified with primers oJLG1872 and oJLG1876.

**Relevant DNA sequences**

Sequence of the $mVenus_{Q69M}$ gene, produced by gene synthesis by Biomatik. The coding sequence is in upper case. The RBS is in lower case.

>mVenusQ69M
taaggaggattttagaATGGTTAGCAAAGGCGAAGAATTATTTACAGGCGTTGTGCCGATTC
TTGTGGAATTAGATGGAGATGTCAACGGCCATAAATTTTCAGTGAGCGGCGAAGGAGAAGGC
GATGCTACATACGGAAAACTTACACTGAAACTGATCTGCACAACAGGCAAACTGCCGGTTCC
GTGGCCGACACTGGTGACAACACTTGGATATGGCTTAATGTGTTTTGCACGCTATCCGGATC
ATATGAAACAACATGATTTCTTTAAATCTGCGATGCCGGAAGGATACGTCCAGGAAAGAACA
ATTTTCTTTAAAGATGATGGCAACTACAAAACACGCGCTGAAGTCAAATTTGAAGGAGATAC
ACTTGTTAACAGAATCGAACTGAAAGGCATCGATTTTAAAGAAGATGGAAACATCCTGGGCC
ATAAACTTGAATACAACTACAACTCACATAACGTTTACATTACAGCTGATAAACAGAAAAAT
GGAATCAAAGCCAACTTTAAAATCCGCCATAACATCGAAGATGGCGGAGTGCAGCTTGCCGA
TCATTATCAACAGAATACACCGATTGGAGATGGCCCGGTCCTGCTTCCGGATAACCATTACC
TGTCTTACCAATCAAAACTTAGCAAAGATCCGAATGAAAAACGCGATCATATGGTCTTACTG
GAATTTGTTACAGCAGCGGGAATCACATTAGGCATGGATGAACTGTATAAATGA

Sequence of the DSM 319 *ponA* gene. The coding sequence is in upper case. The RBS and putative terminator are in lower case. In bold is the cytosine that was replaced by adenine in strain WH320.

>ponA_DSM319
gcgttgaaaggtaggagttttATGTCAGATAATTATCGCTCTCGTGAAGAACGACGAAGAG
CTATGAATGACAACAAACCAGAAGGTAAATCTCAAGGAAAACCGAAAAAGAAGAAAAAAGGA
GGCCTATTTCGTAAAATTGTTGCCGCAGTTTTACTAATAGGAATCATCGGGCTCATTGCCGG
AGTGGGAACTTTTTTTGCTATGATAAGCGATGCTCCAAAAGTTGATGACTCGGTATTAAAAA
ACTCTTTTTCATCTAAAGTATATGCAAACGATGGAAAGACAGTTGTAAAAGAAATTGGAGCT
CAAAAACGTACATATGTGCAATACGATGAAATCCCTCAAGTCGTCAAAAACGCATTTATCGC
TACGGAAGACGTTCGTTTTTATAAACATCACGGCGTCGACTTTTACCGAATTGGCGGAGCAT
TAATGGCGAACTTTAAAAACGGCTTCGGTGCTGAGGGCGGTAGTACGATTACTCAGCAAGTG
GTAAAGAATTCGTTCTTATCACCAAAGAAGACGGTTAAACGTAAAGTACAAGAAATGTGGTT
AGCTTACCAGCTTGAGCGTAAATATTCTAAGCAACAAATTCTTGAAATGTATTTAAACAAAA
TATATTTCGCATCAGGTGAAGTATACGGTATTGAACGAGCAG**C**TGAGAAATTTTACGGAGTT
AAAAGCGTAAAAGATTTAACTCTTGATGAAGCTGCTATGTTGGCAGGCTTACCAAAAGCGCC
AACTACTTATAACCCAGTTACAAACCCAGAGAATGCTACTAAACGCCGTAACACAGTATTAA
ACTTAATGGCTAAAAACGGCTTCATTACACAAGCCGAGGCCGACAAAGCAAAACAAGTAGAT
GTACAGGCTCACTTAGCTAAACAAACTGAATCAACATCTAAGTACGATGTATTTATTGATCA
AGTAATCGAGGAAGTAAAAAAGAAAACAGACGCGGATCCATTTTCTGCGGGCTTAGAAATTT
ATACAACGCTTGACACTAATGCTCAGGACTATGTGGATGATGTGATGAATAACAAAGTAGTC
AACTTCCCGAACGATAAATTCCAAGCAGGTCTTGTCGTTGTAGATAATGAAACAGGCGGTAT
TCAAGCAATTGGAGGCGGACGTAATCGTGTATCAGGCGGCTTAAACTTTGCAACAAATATGA
GACGCCAGCCCGGCTCAACAATTAAGCCGATCTTAGATTACGGTCCAGCGATTGAGAACTTA
AAGTGGTCCACTGGACAACCGCTAAAAGATGAAGAATATCATTATTCAAACGGCACGCCAAT
CCGAAACTTCGGACGCAATTATAAAGGCTGGGTTTCAGCGCGTGAAGCTTTAGGACGCTCAC
TGAACATCCCTGCTTTAAAAGCGTTTCAAGCAACCGGTGCGGACAAAGCAAAAGAGTTCGCT
ACGGGCCTTGGGCTAAAATTAGACGAAGAAGACGGCGAAGCTTATTCTGAATCTTATTCTAT
CGGAGGATTCCGTACAGGTGTTACACCTCTACAAATGGCTGGAGCCTATAGCGCTTTTGCAA
ATGAAGGTATGTACAATGCACCGCATACAGTGACAAAGGTAAAATTCCCTGATGGAACAGAA

ATTGATTTAACACCAAAATCAAAACGCGCAATGCAAGATTACACGGCATTTATGATGACGGA
CATGATGAAATCTGTTGTAAACGAATCATATGGTACAGCTCGTGCTGTTCGTACACCAGGAT
TAGAAATTGCGGGTAAAACGGGAACTTCGAACTTTAGTGCGGCAGAAAAACAACGATACAAC
ATTCCAGACGCTGGTGAAAAAGACGTTTGGTTTAATGGGTTTACGCCAAACTATACGATTTC
CGTATGGACAGGCTATGAAACTGCTAAAGACGGCTATATTTTAAGTCCTAATGAAACAGGCA
TGGCAAAAAATATCTTTAAGCATGTTATTACTCATATTTCTGAAGGCAAACCTAAAGGAAGT
TTCAAGCAGCCGGACAGCGTAGTTGAACGTTCCGTGGAAAAACCAACGGGCAAAGACTTAAA
ACGCCCAAGCGGCTCTACTCCTTCGAGCTTAATTACTAAAGAATACTTTGTACGCGGTACTG
CACCGACAAAAGTATCTCAAACGTTTGTGAAGAAAGAAAAAGAAGAAAAGAAAGAAGAGAAA
GAAGAGAAAAAAGAGGACAAAAAAGAGAAAAAAGACAATAAAAATGTGAAGCCAAAAATTAC
TGCTTCGTATAGCGGAGGAGCAATTTCAGTTAACTGGTCAGCGTCAAACGCAGCCGGTGATA
CGACCTTTACAGTCACAATGTCATCTCCTAACGGTTCTGAACAACTCGCATCTGGCGCAGGA
GCCGGTGGAAAAACAATTTCCAACCCAGCACCAGGCGCAACGTATAGTTTCACCGTAACAAC
AAATACGGGCGAAAGCGCTAGTTCTTCTGTAAAAGTCCCTGATTCATCAGATCAAGCTACCG
AAGATGATTCTACTAAGCCTGAAGATGATGGAACAACGGACGATAATCAAAAGGAAGACAAT
AGCAACACTGATCAAAACAACGGCTCTGATCAAGGTGACGATTCATCTACTCAGCCGGATGA
TAACCAAAACAGCAATAATGGCAGCAATAATGGCAGCAATAACGGCGGTAACAATAGCGGCA
ATACCGATAATAACAATAATGATAGCGGCAATGCTAACGATGGTAACAGCAACAACAACAAT
AATAACGGAGGCACAACAACTCCTCCAGCAGACAACGGTGGAAGTGACTCAGGAAGTCAGAC
ACCACCAACAACCCCTGCTACACCATCGACGCCGTCAACACCTGCTACACCGCAGCAAGAGC
CAACAACTGAAAATTAAcgtagcagtaaaaagcccctacacgtgtaggggctttatttcttg
tgaagaataagca
